# Angiogenesis precedes myogenesis during regeneration following biopsy injury in skeletal muscle

**DOI:** 10.1101/2022.07.23.501245

**Authors:** Nicole L. Jacobsen, Aaron B. Morton, Steven S. Segal

**Affiliations:** Department of Medical Pharmacology and Physiology, University of Missouri; Columbia, MO; Dalton Cardiovascular Research Center; Columbia, MO; Department of Biomedical Sciences, University of Missouri; Columbia, MO; Department of Biomedical, Biological, and Chemical Engineering, University of Missouri; Columbia, MO

**Keywords:** Skeletal muscle, biopsy injury, angiogenesis, myogenesis

## Abstract

**Background:** Acute injury to skeletal muscle damages myofibers and fragments capillaries, impairing contractile function and local perfusion. Myofibers and microvessels regenerate from satellite cells and from surviving microvessel fragments, respectively, to restore intact muscle. However, it is unknown whether myofiber regeneration and microvascular regeneration reflect interdependent processes or may proceed sequentially.

**Methods:** To investigate the temporal relationship between myogenesis and angiogenesis during regeneration, a punch biopsy (diameter, 2 mm) was performed through the center of the gluteus maximus (GM) muscle. Complete removal of all tissue components created a void into which regeneration was evaluated through 21 days post injury (dpi). Confocal imaging and histological analyses of whole-mount GM preparations and GM cross sections assessed the growth of microvessels and myofibers into the wound. Regeneration of perfused microvessels was evaluated *in vivo* by injecting fluorescent dextran into the circulation during intravital imaging.

**Results:** A provisional matrix filled with PDGFRα^+^ and CD45^+^ cells spanned the wound within 1 dpi. Regenerating microvessels advanced into the matrix by 7 dpi. At 10 dpi, sprouting and intussusceptive angiogenesis produced disorganized microvascular networks and spanned the wound with perfusion by 14 dpi. In striking contrast, the wound remained devoid of myofibers at 7 and 10 dpi. Myogenesis into the wound began by 14 dpi with nascent myofibers traversing the wound by 21 dpi. Regenerating myofibers and microvessels were less well organized than in the surrounding (uninjured) muscle.

**Conclusions:** Angiogenesis precedes myogenesis following punch biopsy injury of adult skeletal muscle. Regenerating microvessels encompass the wound and become perfused with blood prior to colocalization with regenerating myofibers. These findings infer that a microvascular supply supports the metabolic demands of regenerating skeletal muscle. Finding that regenerated microvascular networks and myofibers are disorganized within the biopsy site suggests that loss of guidance cues upon complete tissue removal impairs re-establishment of canonical skeletal muscle structure.

## Background

Acute injury to skeletal muscle damages myofibers, triggers an inflammatory response, and fragments capillary networks (1–3). Although the events and time course of the inflammatory cascade (3) and the regeneration of myofibers from resident stem cells [satellite cells (SCs)] following injury are well defined (1, 4–6), data regarding the recovery of the microcirculation following injury are limited. Following exposure to myotoxins, freezing, or physical trauma, capillary blood flow is abolished within 1 day post injury (dpi) but begins to recover by 5 dpi as capillary networks reform (1, 7). Although initially disorganized, newly-formed networks remodel coincident with the maturation of regenerated myofibers by 21 dpi (2, 7, 8).

In accord with their spatial proximity, crosstalk between SCs and capillary endothelial cells (ECs) is integral to the regeneration of intact skeletal muscle and SC self-renewal (9, 10). Moreover, as the immune response transitions from inflammation and degeneration to a milieu that promotes healing (3, 11), SC proliferation is stimulated (12) along with gene co-regulation between angiogenesis and myogenesis (13). However, it is unknown whether microvascular regeneration and myofiber regeneration reflect interdependent processes or may proceed sequentially following injury.

To test whether a temporal relationship exists between myogenesis and angiogenesis during regeneration in adult skeletal muscle, we developed a model of complete tissue removal using punch biopsy of the mouse gluteus maximus (GM) muscle. We hypothesized that following removal of all tissue components, angiogenesis precedes the formation of new myofibers to support the metabolic demands of regenerating skeletal muscle. We report here that creating a void into which regeneration proceeds reveals that angiogenesis occurs prior to myogenesis after biopsy injury. These findings establish a foundation for detailed studies of distinct versus shared regulation between regeneration of myofibers and their vascular supply.

## Methods

### Animal care and use

Male and female mice (C57BL/6, Jackson Laboratory; Bar Harbor, ME) were bred and housed in animal care facilities of the University of Missouri. Mice (weight, ~30 g) were studied the age of ~4 months. In reporter mice bred on a C57BL/6 background [Cdh5-Cre^ERT2^ (14) × ROSA26^mTmG^ (#007676, Jackson Laboratory)], Cre recombination for expression of membrane bound green fluorescent protein (GFP) in ECs was induced through intraperitoneal injection of 100 uL tamoxifen (10 mg/mL + 5% ethanol in corn oil; #T5648, Sigma-Aldrich; St. Louis, MO) on 3 consecutive days; at least 1 week elapsed after the first injection prior to study. All mice were maintained under a 12:12 h light/dark cycle at 22-24 °C with fresh food and water *ad libitum*. To avoid any order effect, collection of data at criterion timepoints was randomized. Prior to muscle injury, intravital microscopy, or tissue collection, a mouse was anesthetized [ketamine (100 mg/kg) + xylazine (10 mg/kg) in sterile saline; intraperitoneal injection]. Mice were euthanized at the end of an experiment by anesthetic overdose and cervical dislocation.

### Punch biopsy

A mouse was anesthetized and positioned on an aluminum warming plate to maintain body temperature at 37 °C. As needed, supplemental injections of anesthesia (~20% of initial) were given to maintain a stable plane of anesthesia confirmed by lack of withdrawal to a toe pinch (monitored every 15 min). Skin covering the left GM was shaved and sterilized by swabbing 3× with Betadine and 70% alcohol. While viewing through a stereomicroscope, the mouse was positioned on its abdomen and a ~1 cm incision was made through the skin overlying the GM near the spine. The skin was retracted to expose the GM and irrigated with sterile saline. A hole was made through the center of the GM with a sterile 2 mm diameter biopsy punch (MediChoice #DP0200, Owens & Minor; Mechanicsville, VA); this size is below the limits of muscle loss for effective regeneration (15) while large enough to create a void into which regeneration could be reliably studied. Anatomical landmarks provided a reference for consistency in the site of biopsy. The skin was closed with sterile 6-0 nylon sutures (UNIFY #S-N618R13, AD Surgical; Sunnyvale, CA). This entire procedure required ~20 min. For recovery, the mouse was placed on a heated platform, monitored until consciousness and ambulation were restored (~2-3 h), then returned to its original cage. Normal activity and behavior were routinely observed within 24 h. Regeneration of tissue components into the wound was evaluated at key timepoints through 21 days post injury (dpi) with uninjured mice (0 dpi) as controls.

To evaluate cellular damage surrounding the biopsy site (Fig S1), Evans Blue dye (EBD) (1% solution in sterile saline; #E2129, Sigma) was injected intraperitoneally (10 μL/g body mass) following the surgical procedure. At 1 dpi, the GM was dissected (as below) to image EBD. Images were acquired with a 4× objective on an E800 microscope coupled to DS-Fi3 camera using Elements software (all from Nikon; Tokyo, Japan).

### Dissection of the gluteus maximus muscle

A mouse was anesthetized, the surgical area was shaved, the mouse was placed on a warming plate, and the skin overlying the GM was removed with scissors. Exposed tissue was continuously superfused with a bicarbonate-buffered physiological salt solution (bbPSS; pH 7.4, 34-35°C) containing (in mM) 131.9 NaCl_2_ (Fisher Scientific; Pittsburg, PA), 4.7 KCl (Fisher), 2 CaCl_2_ (Sigma), 1.17 MgSO_4_ (Sigma), and 18 NaHCO_3_ (Sigma) equilibrated with 5% CO_2_/95% N_2_. While viewing through a stereomicroscope, excess connective tissue was removed using microdissection while avoiding the site of injury. The GM was cut from its origin along the lumbar fascia, sacrum, and iliac crest then reflected away from the body to view its vascular supply from the ventral surface (7, 16).

### Intravital microscopy

The exteriorized GM was spread onto a transparent rubber pedestal (Sylgard 184; Dow Corning; Midland, MI) and pinned at its edges to approximate *in situ* dimensions. Exposed tissue on the body of the mouse was covered with plastic film (Glad Press n’ Seal) to prevent dehydration, the preparation was transferred to the stage of a Nikon E600FN microscope, and the GM equilibrated for 30 min while continuously superfused with bbPSS at 3 mL/min. Supplemental doses of anesthetic were given throughout the experimental protocol (duration, 2-3 h) to maintain a stable plane of anesthesia (as above). To assess vascular perfusion, fluorescein isothiocyanate (FITC) conjugated dextran (70 kDa; 10 mg/mL sterile saline) was injected (200 μL) into the systemic circulation via the retroorbital sinus and allowed to circulate for ~10 min. The GM was illuminated with a mercury lamp for fluorescence imaging using a filter cube appropriate for circulating FITC or transgenic eGFP. Images were acquired through Nikon Plan Fluor 4×/0.13 or Plan Fluor 10×/0.3 objectives coupled to a low light CMOS FP-Lucy camera [Stanford Photonics, Inc.; Palo Alto, CA (SPI)] and displayed on a digital monitor. Time lapse images were recorded at 40 frames/s using Piper Control software (SPI).

### Confocal imaging of fresh whole mount muscle preparations

The GM was removed from a Cdh5-mTmG mouse and placed in a custom imaging chamber with the ventral surface facing the objective. To optimize resolution of the microvasculature, a drop of PBS (~10 μL) was added to the chamber and the GM was flattened by placing a glass block (2 cm × 2.5 cm × 1 cm; mass, 7.8 g) on the dorsal surface. Images were acquired with a HC PL APO 10/0.40 CS2 objective on an inverted laser scanning confocal microscope (TCS SP8) using LASX software (all from, Leica Microsystems; Buffalo Grove, IL). A Dragonfly High Speed Confocal Microscope System [Leica DMi8 microscope coupled to an Andor Dragonfly 200 and integrated Fusion software (Oxford Instruments; Abingdon, United Kingdom) was also used to acquire images. To image the entire wound, tile scans (3 × 3 grid encompassing ~3.5 mm × 3.5 mm) were acquired with a Leica HC PL APO 10×/0.45 objective and stitched using integrated software. Using maximum projection z-stacks acquired with a Leica HC PL IRAPO 20×/0.75 objective, 10 random microvessels were chosen within a region of interest (ROI; 493 × 493 μm). Diameters of regenerating microvessels were measured at the midpoint between two branch points and averaged for a given GM; 5-7 GM were analyzed at respective dpi.

### Immunostaining

An excised GM was immobilized by pinning the edges in the well of a 12-well plate coated with Sylgard 184. After washing with PBS, a region of muscle (~5 mm × 5 mm) containing the injury surrounded by undamaged tissue was trimmed and prepared for either whole mount immunostaining or frozen in Tissue Tek OCT compound (VWR International LLC; Radnor, PA) to obtain tissue cross sections for histology.

Whole mount preparations were fixed in 2% paraformaldehyde for 30 min, washed in PBS, and placed in blocking buffer (2% bovine serum albumin, 4% normal donkey serum, 0.5% triton X-100 in PBS) for 30 min. Preparations were then incubated overnight at 4 °C with validated primary antibodies: anti-rat CD31 [platelet endothelial cell adhesion molecule, [PECAM-1; 1:400, #550274, BD Pharmingen; San Diego, CA; (17)] to identify endothelial cells; anti-rabbit CD45 [leukocyte common antigen; 1:200, #ab10558, Abcam; Cambridge, UK; (18)] to identify inflammatory cells; and anti-goat PDGFRα [1:200, #AF1062, R&D Systems; Minneapolis, MN; (19)] to identify fibroadipogenic precursor cells (FAPs) (20–22). Preparations were washed in blocking buffer, incubated with secondary antibodies for 30 min, washed again, then rinsed in PBS before transfer to an imaging chamber for confocal image acquisition (as above).

For tissue cross sections, trimmed and washed GM samples were transferred to a cryomold containing OCT compound with black silk suture (2-0 thickness; length, 2 mm) positioned adjacent to the injury for spatial reference. The cryomold was frozen in isopentane cooled in liquid nitrogen and stored at −80 °C until sectioning. Cross sections of frozen GM were cut (thickness, 10 μm) with a cryostat (HM 550 Cryostat, Thermo Scientific; Waltham, MA) at the center of the injury referencing to the silk suture.

For fluorescence imaging, sections were fixed in 4% paraformaldehyde for 10 min and then stained with CD31 (1:500, BD Pharmingen), myosin heavy chain [1:5, MF-20, Developmental Studies Hybridoma Bank, The University of Iowa Department of Biology; Iowa City, IA; (23)], laminin [(1:200, #PA1-16730, Invitrogen; Waltham, MA; (24)], and appropriate secondary antibodies (1:400, AlexaFluor, Fisher). Prolong Gold containing DAPI (Fisher) was added before slides were coverslipped. Sections were imaged using appropriate filters on a Nikon E800 microscope with a DS-Qi2 camera and Nikon Elements Software.

The distance devoid of myofibers (e.g., diameter of remaining visible wound) was measured along GM cross sections accounting for curvatures within the specimen (Fig. 4). Within this region, the total area of CD31^+^ staining was measured to evaluate microvascular density within the wound preceding myofiber ingrowth.

### Histochemistry

To visualize collagen deposited within the provisional matrix in whole mount preparations, the GM was removed, permeabilized in 0.5% Triton X-100 in PBS, and incubated for 1 h with PicroSirius Red (1% Direct Red in saturated picric acid). Thereafter, GM were treated for 30 min with 0.5% acetic acid in ddH_2_O and rinsed in 100% EtOH. Images were acquired as described for EBD.

### Statistics

Summary data are displayed for individual GM preparations along with means ± s.e.m. Statistical analyses were performed using Prism 9 software (GraphPad Software, Inc.; La Jolla, CA). One-way ANOVA with Tukey’s post hoc tests were used to determine statistical significance between time points with P < 0.05 considered significant. Values for ‘n’ refer to the number of GM analyzed; one GM was studied per mouse.

## Results

### Punch biopsy creates a void into which regeneration advances

Muscle injury was created using a biopsy punch (diameter, 2 mm) to remove a volume of muscle below the critical threshold for regeneration [Fig. 1A; (15)]. A circular hole through the center of the GM resulted in minimal collateral damage to the surrounding tissue as evidenced by nominal uptake of EBD into the ends of myofibers bordering the wound at 1 dpi (Supp. Fig. 1). Observed in anesthetized mice, intravascular injection of FITC dextran confirmed the perfusion void within the wound while blood flow was preserved to tissue surrounding the injury (Fig. 1B). Sirius red staining of whole mount GM preparations revealed the deposition of collagen to create a provisional matrix (25) that nearly spanned the wound at 1 dpi (Fig. 1C); immunostaining demonstrated invasion of this matrix by PDGFRα^+^ cells (fibroadipogenic progenitor cells, FAPs) accompanied by CD45^+^ (inflammatory) cells (Fig. 1D).

**Figure 1.**
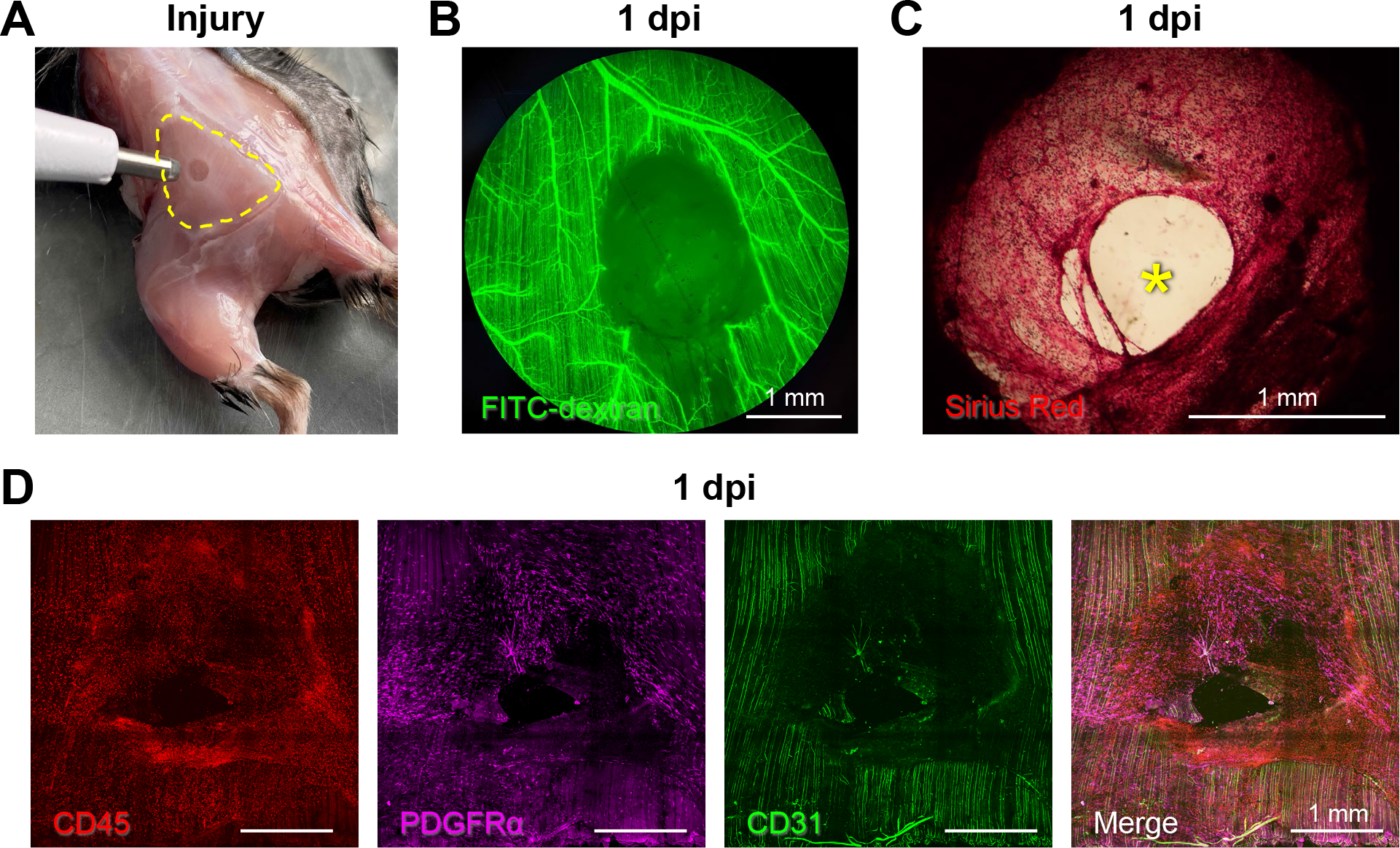
Punch biopsy removes all skeletal muscle tissue components. **A.** A biopsy punch was used to create a local injury (diameter, 2 mm) through the GM (muscle perimeter outlined by yellow broken line). **B.** Microvascular perfusion was disrupted at the site of the injury at 1 dpi. Intravascular injection of fluorescent dextran showed the empty biopsy site in addition to a small unperfused region downstream of the injury. **C.** Representative Sirius red staining of collagen at 1 dpi highlighting deposition of a provisional matrix in the wound; * indicates residual gap. **D.** Stitched tile scan images of immunostained whole mount GM at 1 dpi showing masses of inflammatory cells (CD45^+^, red) and fibroblasts (PDGFRα^+^, magenta) invading the provisional matrix; capillaries (CD31^+^, green) retain parallel orientation along surrounding uninjured myofibers.

### Microvascular growth into the wound

Using an EC-specific Cre driver [Cdh5-CreERT2; (14)] and tamoxifen-induced recombination of the Rosa26 mTmG locus to genetically label the endothelium with membrane-bound eGFP (Cdh5-mTmG mice), we evaluated microvascular regeneration following punch biopsy. Angiogenic sprouts reflecting endothelial tip cells (26) were apparent at 5 dpi (not shown) and extended into the wound at 7 dpi (Fig. 2A&B). Nascent microvessels also underwent internal division (i.e., intussusceptive angiogenesis) as networks expanded to extend across the wound (Fig. 2C-E). Through 14 dpi, microvessels split into daughter vessels generating numerous branches with short interconnections. The angiogenic response from 7-14 dpi progressed as a centripetal gradient of microvessel growth from the edges towards the center of the wound.

**Figure 2.**
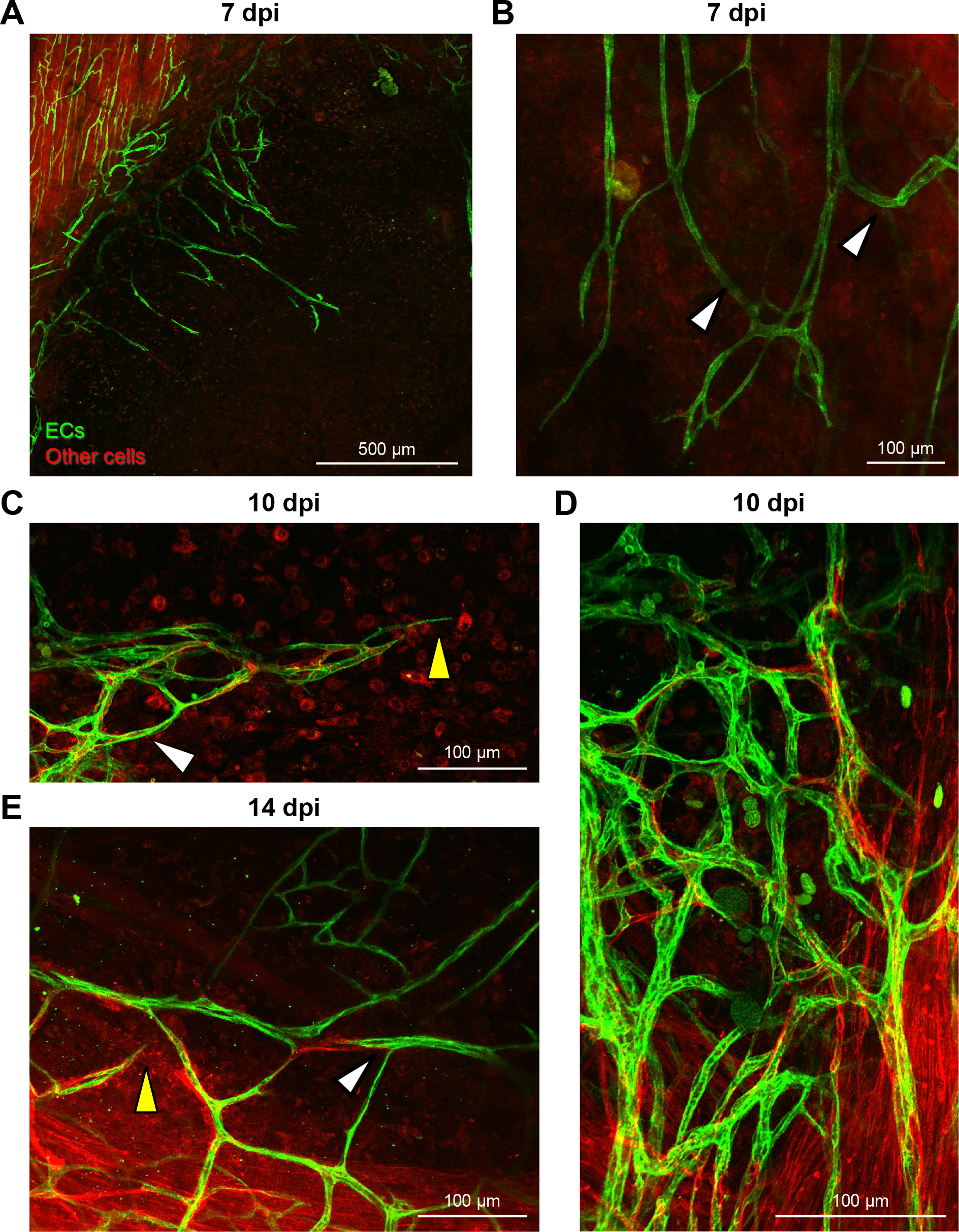
The microcirculation regenerates through sprouting and intussusceptive angiogenesis. **A.** At 7 dpi, nascent microvessels emigrate from existing vasculature at wound edge. **B.** At 7 dpi, microvessels are nonuniformly enlarged (arrowheads) and extend into the wound with random orientation. **C.** Intussusceptive angiogenesis (white arrowhead) at 10 dpi is integral to revascularization while sprouting continues (yellow arrowhead). **D.** Sites of robust angiogenic activity at 10 dpi create irregular microvascular plexuses. **E.** At 14 dpi, sprouting (yellow arrowhead) and intussusceptive angiogenesis (white arrowhead) expand regenerating microvascular networks. Images are from Cdh5-mTmG reporter mice in which ECs are green (eGFP) and other cells are red.

By 14 dpi, microvascular networks spanning the wound were perfused with blood but lacked hierarchical structure or definitive patterns of blood flow (Fig. 3, Supp. Fig. 2). Instead, nascent microvessels were clustered into zones of robust angiogenic activity with random orientation (Fig. 2D). In addition, microvessel diameter significantly increased from 4.1 ± 0.3 μm in uninjured muscle to 7.4 ± 0.4 μm at 7 dpi and 9.0 ± 0.6 μm at 14 dpi (Fig. 3D). Intravital microscopy revealed that only the microvessels having the largest diameter were continuously perfused; most segments contained stationary red blood cells or plasma alone at 14 dpi (Supp. Fig. 2). In perfused microvessels, the direction of blood flow oscillated and was interspersed with periods of flow cessation.

**Figure 3.**
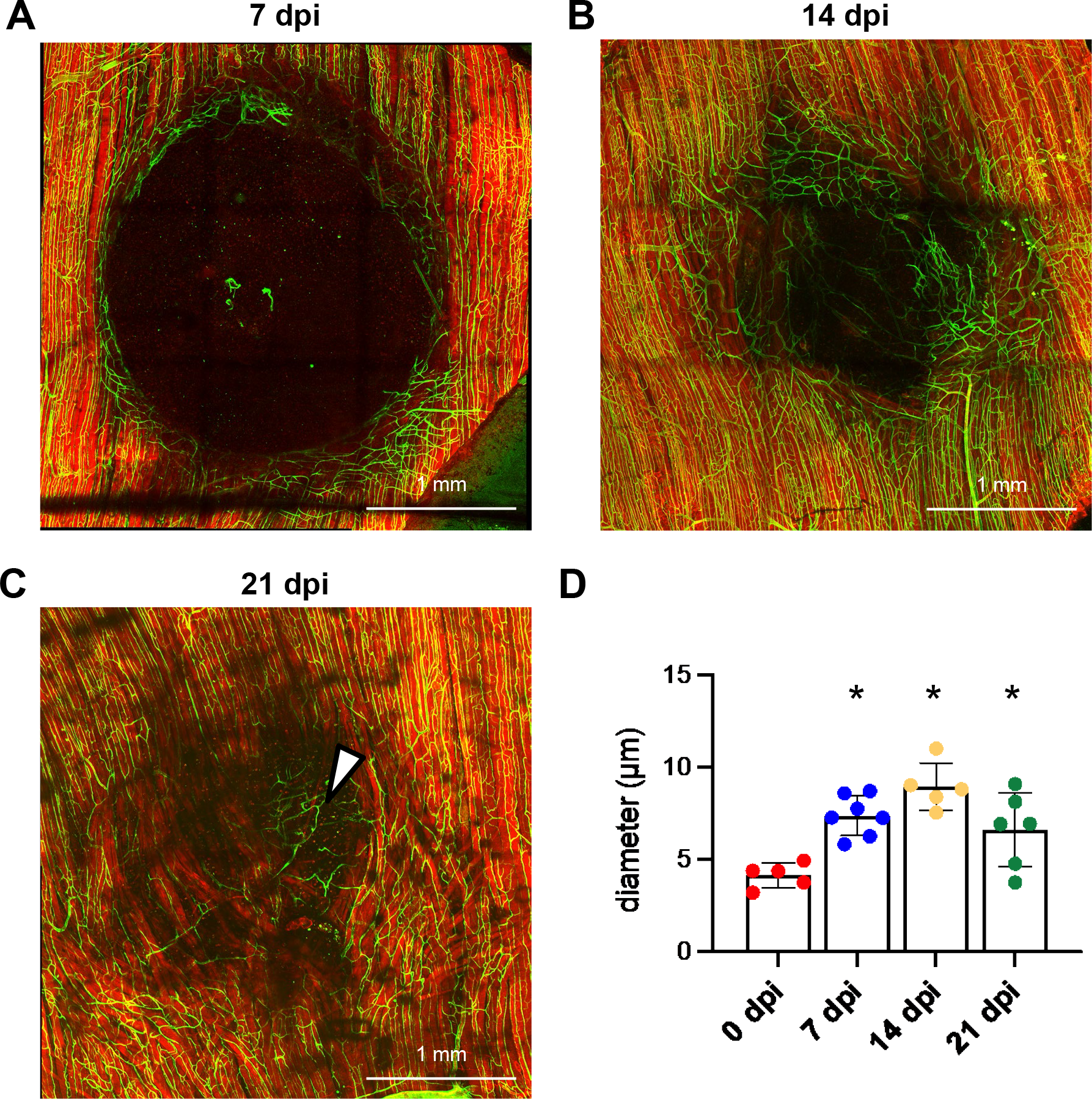
Tile scanned images showing time course of microvascular and myofiber regeneration after punch biopsy. **A.** At 7 dpi, angiogenesis begins around edges of wound. **B.** At 14 dpi, regenerating microvessels have spanned the wound which remains devoid of myofibers **C.** At 21 dpi, myofibers traverse the wound while capillary networks are disorganized. Dark areas within the regenerated GM reflect interweaving among adjacent myofibers in contrast to parallel (flat) organization of surrounding (uninjured) myofibers. Within the wound, microvessels remodel (arrowhead) to supply a cluster of adipocytes (not visible) where myofibers have not formed. Images are from Cdh5- mTmG mice in which ECs are green (eGFP expression) and all other cells are red (tdTomato expression). **D.** Summary data of microvessel diameters at criterion time points; n = 5-7 mice per time point, *P < 0.05 versus 0 dpi.

By 21 dpi, extensive microvascular networks encompassed the site of injury (Fig. 3C) with diameters (6.6 ± 0.8 μm) returning towards control. Nevertheless, the parallel arrangement of microvessels observed in the surrounding uninjured muscle was not restored. Instead, numerous anastomoses created circuitous pathways of local perfusion.

### Regeneration of myofibers into the wound

In striking contrast to the time course of revascularization, the wound remained devoid of myofibers at 7 dpi, a time at which myofiber regeneration would be well advanced in other models of skeletal muscle injury (1, 7). As myofibers began to penetrate the provisional matrix and regenerate into the wound at 10-14 dpi (Fig. 3), they were densely packed, interwoven, and coursed along the wound edge before integrating with undamaged myofibers. Additionally, regenerating myofibers were smaller in width (i.e., cross sectional area) than in surrounding (uninjured) tissue, consistent with other muscle injury models (1, 5, 7, 27). Newly-formed myofibers intertwined as they spanned the wound at 21 dpi. In some preparations, clusters of adipocytes occupied space where myofibers had not regrown (Fig. 3C, Supp. Fig. 3). In turn, regenerating capillary networks remodeled in accord with adipocyte morphology (Fig. 3C).

To confirm that revascularization occurred prior to myofiber regeneration, GM cross sections (thickness, 10 μm) were immunolabeled for CD31 to identify ECs, myosin heavy chain (MyHC) for contractile myofibers, laminin to detect basal laminae, and DAPI for nuclei. Congruent with data from intravital and confocal imaging, healing progressed into the wound from 7 dpi through 21 dpi with regeneration of microvessels preceding that of myofibers (Fig. 4A-C). The length of the injured region devoid of myofibers (corresponding to the diameter of existing wound) was used as an index of healing over time. This length was 1512 ± 153 μm at 7 dpi which progressively decreased to 475 ± 134 μm at 14 dpi and 71 ± 71 at 21 dpi (Fig. 4D). We quantified the area occupied by ECs within the region lacking myofibers as an indicator of microvascular density that preceded myogenesis. This EC area was 2794 ± 1126 μm^2^ at 7 dpi, 3366 ± 930 μm^2^ at 14 dpi, and decreased to 700 ± 700 μm^2^ at 21 dpi (Fig. 4E), reflecting the growth of myofibers into the wound, colocalizing with ECs (microvessels) as regeneration advanced. In contrast to centrally-located nuclei identifying regenerated myofibers (1, 4, 5), adjacent fibers within the injury often had nuclei localized to their periphery (Fig. 4F), suggesting that the removal of stroma affects localization of myonuclei during myogenesis post injury.

**Figure 4.**
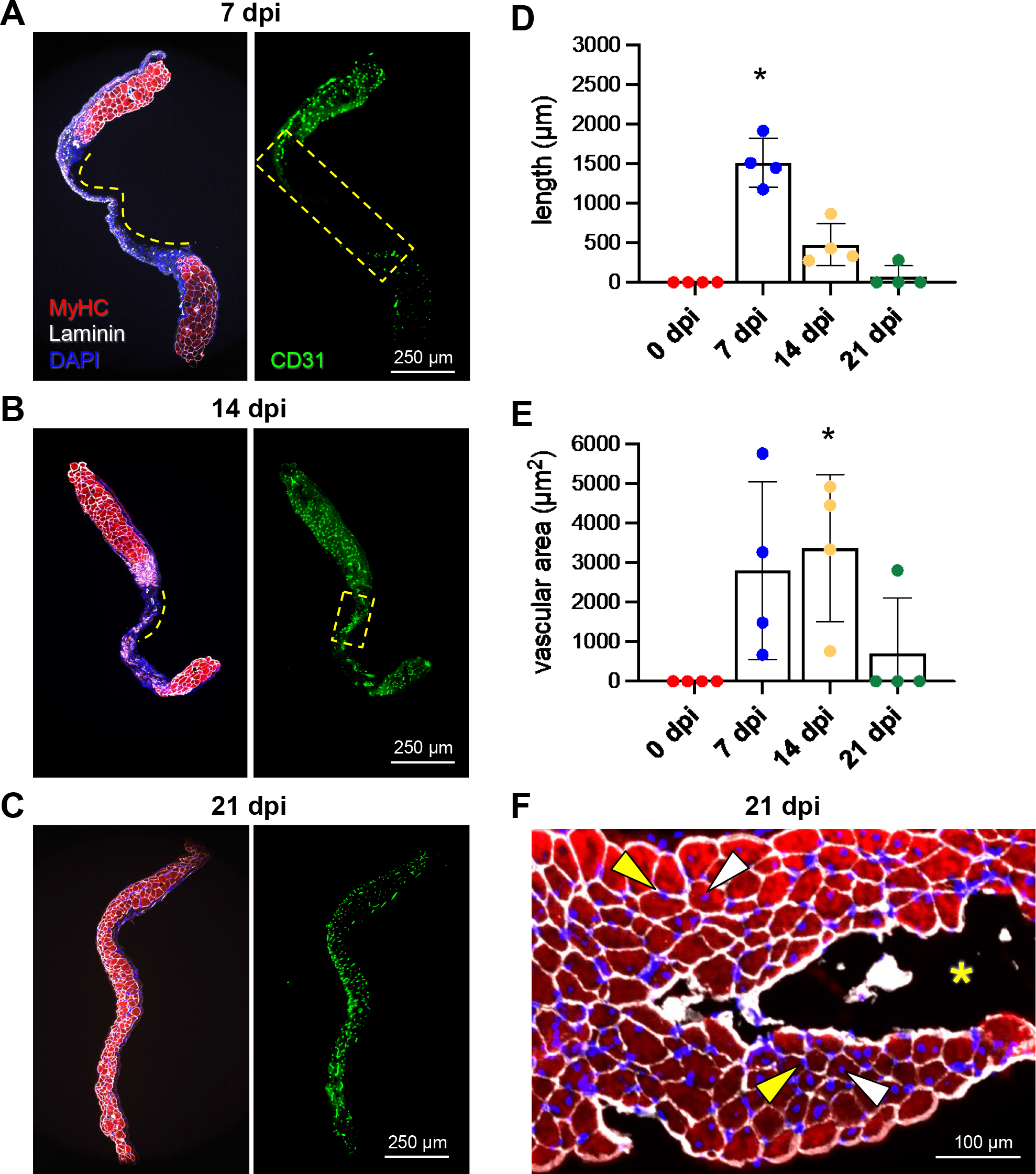
Angiogenesis precedes myogenesis during regeneration. Representative images of GM cross sections from the center of the wound at **A.** 7 dpi, **B.** 14 dpi, and **C.** 21 dpi. MyHC (myofibers): red; laminin (basal laminae): white; CD31 (ECs): green; nuclei: blue. **D.** Summary data for length of region devoid of myofibers (shown by broken yellow lines in A and B) and **E.** Vascular area within the wound region devoid of myofibers (outlined by broken yellow boxes in A and B). n = 4 per time point, *P < 0.05 vs 0 dpi. **F.** Cross section of from regenerated region within the wound at 21 dpi stained for MyHC (red) and myonuclei (blue, DAPI). Some myofibers exhibit centrally located nuclei (white arrowheads) while others exhibit peripheral nuclei (yellow arrowheads). Yellow asterisk: region within punch biopsy where myofibers did not regenerate.

## Discussion

Punch biopsy is an acute injury that is shown here to dissociate the time course of angiogenesis from that of myogenesis in regenerating skeletal muscle for the first time. In contrast to myotoxins, freezing, or physical trauma, where efferocytosis removes cellular debris while leaving a residual extracellular matrix [ECM (28–31)], punch biopsy removes all tissue components to create an empty void. In the GM of adult male and female mice, creating such a void through the center of the muscle resulted in the formation of a provisional matrix (25) filled with immune cells and FAPs at 1 dpi. Endothelial cells proliferate and migrate from the edges of the wound into the provisional matrix by 7 dpi; a combination of sprouting and intussusceptive angiogenesis resulted in the development of microvascular networks that spanned the wound and became perfused with blood by 14 dpi. Myofibers grew into the wound by 14 dpi and filled the wound at 21 dpi, approximately 1 week after comparable events in the regenerating microvasculature. Microvessels and myofibers within the wound are disorganized compared to uninjured muscle suggesting that guidance cues for regeneration may be lost upon complete tissue removal that would otherwise promote restoration of intact muscle.

### Microvessel and myofiber regeneration following injury

The GM is a thin muscle comprised of mixed fiber type (32) rendering it well-suited for studying microvascular networks in the mouse (16). Previous studies evaluating the GM microcirculation following injury with BaCl_2_ show that the microcirculation regenerates and recovers function concomitant with myofibers through 21-35 dpi (2, 7, 8). These findings in the GM are consistent with injuries to other rodent hindlimb muscles (1, 4, 5, 13, 27). Myofibers and microvessels regenerate *in situ* from SCs and from surviving capillary fragments (i.e., ECs), respectively, to restore intact muscle (1, 2, 4, 7). Myofiber regeneration ensues as the milieu transitions from proinflammatory to one that promotes healing (3, 33). Earlier studies reported that capillarization is governed by the size and growth of regenerating myofibers (34), with an upregulation of angiogenic gene signatures during the early stage of muscle regeneration (35). Supporting this view were findings that ECs enhanced SC growth through secretion of an array of growth factors [VEGF, IGF-1, HGF, bFGF, PDGF-BB (9)]. Complementary studies found that SCs modulate angiogenesis through hypoxia-inducible factor-1 (HIF-1), a transcriptional regulator of VEGF gene expression (36). With SCs remaining in juxtavascular locations, differentiating myogenic cells were both proangiogenic and spatiotemporally associated with angiogenesis (9).

Sensing attractive and repulsive signals in the microenvironment, ECs invade the provisional matrix and began to revascularize the wound at 7 dpi (Fig. 2). With sprouting angiogenesis, endothelial tip cells guide the formation of new capillaries while stalk cells proliferate behind the tip cells to extend the newly forming microvessels (26). Intussusceptive angiogenesis further contributes to microvascular proliferation within the wound (Fig. 2) by forming pillars within the lumen of a vessel that extend to split a preexisting vessel into two daughter segments and modify branch angles at bifurcations (37). During regeneration following hindlimb ischemia in mice, intussusception was the dominant mechanism of angiogenesis (38). The findings presented herein illustrate that both sprouting and intussusceptive angiogenesis contribute to microvascular regeneration following punch biopsy injury. The formation of new microvessels through vasculogenesis (39) may also contribute to revascularization but was not resolved in the present experiments.

### Modulation of SC-EC crosstalk by immune cells and FAPs

Crosstalk between surviving SCs and ECs occurs under the influence of immune cells and FAPs (3, 6, 9–13, 20, 40, 41). FAPs are identified by the expression of mesenchymal stem cell surface proteins (e.g., Sca-1, CD34, PDGFRα) and reside in the interstitial space adjacent to the microvasculature (21, 22, 42). Upon injury, FAPs enter the cell cycle to proliferate, localize to the site of tissue damage, differentiate into fibroblasts and adipocytes [but not into SCs or myofibers (20, 22, 40)], and work in concert with macrophages and ECs to promote myofiber regeneration (6, 22, 41, 43). Our finding that CD45^+^ and PDGFRα^+^ cells invade the provisional matrix within 1 dpi following punch biopsy (Fig. 1) is consistent with earlier studies (13, 20, 22, 43). Activated FAPs deposit ECM proteins (e.g., collagen), secrete myogenic differentiation factors, and recruit CD45^+^ immune cells via paracrine signaling (20, 43). The ensuing cascade of granulocytes, monocytes, and macrophages produce cytokines and chemokines aimed at clearing cellular debris, promoting self-renewal of SCs, and preventing premature myogenic cell differentiation (3, 44).

The ECM aids in cell adhesion, cell-to-cell communication, and differentiation (28, 45). In addition to providing matrix structure, collagen fibers guide the growth of capillaries during regeneration (46). In such manner, the microenvironment of the provisional matrix established by FAPs and immune cells serves as the foundation for the regeneration of intact skeletal muscle (25). However, if the injury exceeds a critical threshold of muscle loss [e.g., a 3-mm diameter biopsy in the mouse quadriceps (15)], inflammation and fibrosis persist. Furthermore, if myoblast migration and fusion are delayed during regeneration, FAPs may also contribute to myosteatosis (42).

### Novel features of punch biopsy injury

Previous studies of muscle injury have not resolved whether regeneration of myofibers and microvessels following injury are interdependent processes or whether angiogenesis and myogenesis proceed as sequential events. Indeed, earlier findings have led to the conclusion that angiogenesis and matrix remodeling occur during the last phase of myogenesis and growth of new myofibers (5, 33). However, data presented herein show that matrix generation and angiogenesis occur first. Such ambiguity may be attributed to the persistence of basal laminae following efferocytosis, an outcome that prevails in established models (1, 4, 5, 11, 27). Preexisting basal laminae can provide guidance during regeneration of microvessels and myofibers (28–31, 46, 47), but a biopsy punch removes the entire tissue. A key finding here is that, in contrast to the concomitant regeneration of microvessels and myofibers discussed above, the onset of microvascular growth into the wound and perfusion across the wound precedes that of myofibers by approximately 1 week. We attribute this difference from other injury models to the removal of the entire tissue, with loss of primed progenitor cells and guidance cues that would otherwise be retained in the residual stroma.

In addition to distinguishing between the time course of angiogenesis versus myogenesis, key differences emerged during regeneration following punch biopsy when compared to other models of muscle injury. In contrast to the onset of sprouting angiogenesis 2-3 dpi following exposure to myotoxins (1, 2, 7), EC sprouts were not apparent until ~1 week following punch biopsy (Fig. 2A). This longer delay may be attributable to prioritized healing around the edges of the open wound prior to EC invasion of the provisional matrix. A delay in angiogenesis (14 dpi) was also observed following freezing (1)], which kills all cells within the injured region.

Nascent microvessels were nonuniformly enlarged through 21 dpi compared to capillaries supplying healthy myofibers in the surrounding tissue. This finding is inconsistent with data from the GM following BaCl_2_ injury, where vessel diameter returned to baseline values and the structure of capillary networks had nearly recovered at this time (8). The irregular blood flow patterns observed within the wound (Supp. Fig. 2) can be attributed to the absence of hierarchical organization of newly-formed networks or established arterio-venous pressure gradients (48), reflecting the absence of structural and functional cues that may otherwise be provided by the residual ECM.

Whereas centrally located myonuclei are a hallmark of regenerated myofibers (1, 4, 5), including the GM following BaCl_2_ injury (7), regenerating myofibers filling the void contained myonuclei positioned centrally in some myofibers while positioned at the periphery of other myofibers, often located adjacent to each other (Fig. 4F). This difference in nuclear localization implies a modified mechanism of myogenesis when compared to other models of skeletal muscle injury and myofiber regeneration (49), which may reflect a distinct microenvironment for regeneration after biopsy injury. That regenerating myofibers intertwine instead of exhibiting their characteristic parallel alignment further indicates that the provisional matrix lacks the structural basis to dictate the canonical arrangement of myofibers (29, 30).

### Summary and Conclusions

The present study demonstrates that creating a hole through the GM muscle enabled definitive temporal resolution of angiogenesis versus myogenesis for the first time following injury in adult skeletal muscle. Following punch biopsy injury to remove all tissue components, a provisional matrix invested with inflammatory cells and fibroblasts is deposited within 1 dpi. Sprouting and intussusceptive angiogenesis revascularize the wound at 7-14 dpi, a time frame that precedes the onset and completion of myofiber regeneration by ~ 1 week. Consistent with other models of injury, myofibers invested with a microvascular supply are restored by 21 dpi. However, regenerated capillary networks and myofibers within the site of injury remain disorganized relative to uninjured muscle, suggesting that the total loss of skeletal muscle ultrastructure disrupts the morphology of regenerating tissue components.

These experiments are the first to illustrate that revascularization precedes the replacement of myofibers in adult skeletal muscle following injury. Our findings are consistent with the interpretation that establishing a vascular supply in the provisional matrix generates conditions that support the regeneration of myofibers (6). The outcome of this study contrasts with injuries in which the basal laminae (and progenitor cells) persist and angiogenesis occurs concomitant with myogenesis (9, 13, 47). The present findings thereby provide a foundation for selectively manipulating variables known to affect respective components of regeneration to restore the functional integrity of skeletal muscle.

## List of abbreviations

Cdh5-mTmG: Cdh5-Cre^ERT2^ × Rosa^mTmG^ transgenic mice;
dpi: days post injury;
EBD: Evans Blue dye;
ECs: endothelial cells;
ECM: extracellular matrix;
GFP: green fluorescent protein;
FAPs: fibroadipogenic precursor cells;
FITC: fluorescein isothiocyanate;
GM: gluteus maximus;
PDGFRα: platelet derived growth factor receptor alpha;
MyHC: myosin heavy chain;
SCs: satellite cells.

## Declarations

### Ethical approval

All procedures were approved by the Animal Care and Use Committee at the University of Missouri (protocol #17720). Experiments were performed in accordance with the National Research Council’s Guide for the Care and Use of Laboratory Animals.

### Consent for publication

Not applicable.

### Availability of data and materials

The data generated and analyzed during the current study are available from the corresponding author on reasonable request.

## Acknowledgments

Yuki Yang contributed to preliminary experiments. Drs. Erika Boerman and Scott Zawieja provided use of their confocal microscopes and imaging processing software. Drs. DDW Cornelison and Charles Norton contributed thoughtful discussion throughout data collection and analysis.

## Author contributions

Conceptualization: NLJ, ABM, SSS; Methodology: NLJ, ABM, SSS; Investigation: NLJ, ABM, SSS; Visualization: NLJ, ABM; Funding acquisition: NLJ, ABM, SSS; Project administration: SSS; Supervision: SSS; Writing – original draft: NLJ; Writing – review & editing: NLJ, ABM, SSS

---

**Supp. Figure 1.**
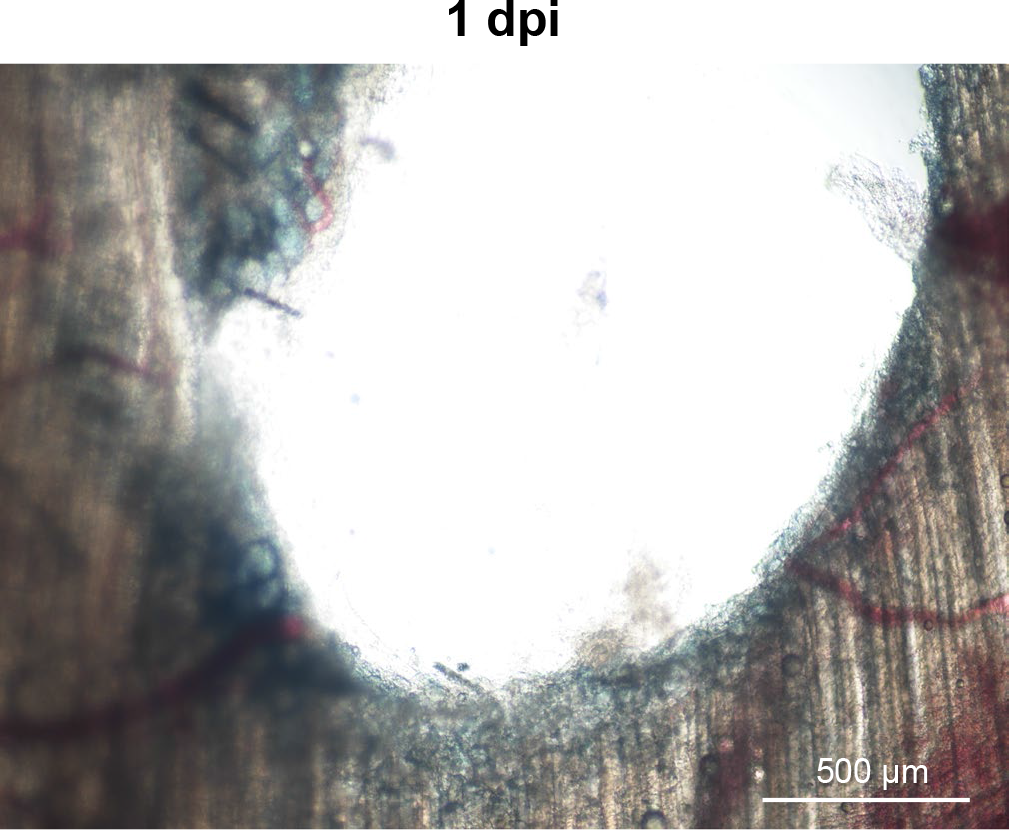
At 1 dpi, Evans Blue dye uptake is restricted to the edges of biopsied myofibers.

**Supp. Figure 2.**
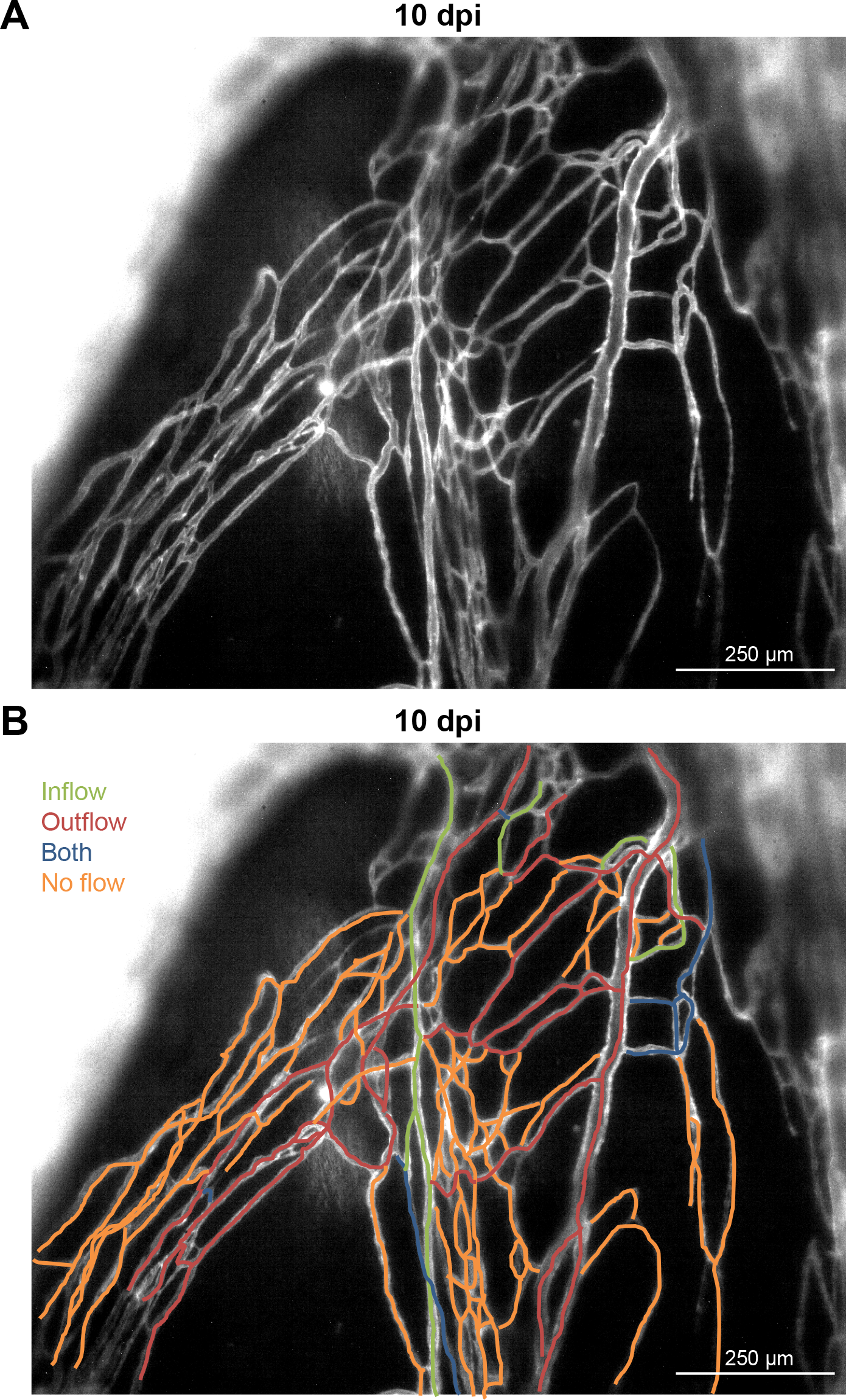
Blood flow within the regenerating wound at 10 dpi was visualized by red blood cell transit during intravital microscopy. **A.** Monochrome image of regenerating microvessels in Cdh5- mTmG mice. **B.** Color coding denotes the presence and direction of red blood cell flow at 10 dpi.

**Supp. Figure 3.**
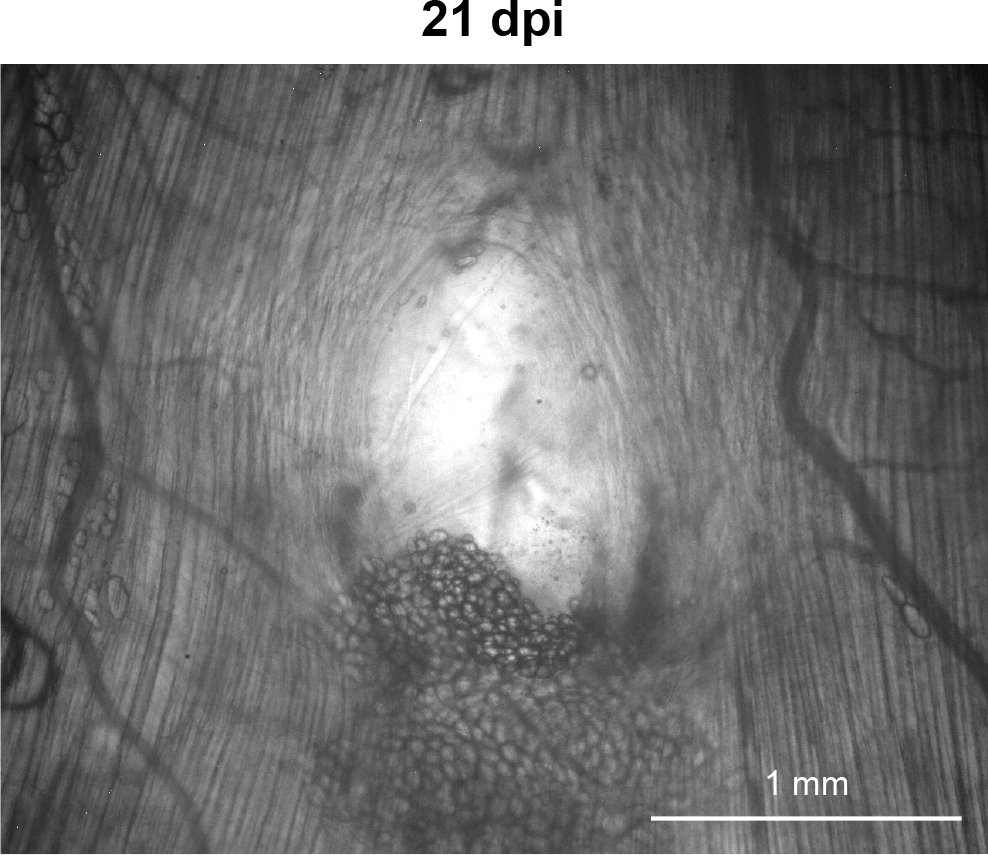
Adipocytes accumulate in the biopsy void if myofibers fail to regenerate at 21 dpi.

